# Alternative transcription cycle for bacterial RNA polymerase

**DOI:** 10.1101/663534

**Authors:** Timothy T. Harden, Karina S. Herlambang, Mathew Chamberlain, Jean-Benoît Lalanne, Christopher D. Wells, Gene-Wei Li, Robert Landick, Ann Hochschild, Jane Kondev, Jeff Gelles

## Abstract

RNA polymerases (RNAPs) transcribe genes through a cycle of recruitment to promoter DNA, initiation, elongation, and termination. After termination, RNAP is thought to initiate the next round of transcription by detaching from DNA and rebinding a new promoter. We used single-molecule fluorescence microscopy to observe individual RNAP molecules after transcript release at a terminator. Following termination, RNAP almost always remained bound to DNA and sometimes exhibited one-dimensional sliding over thousands of basepairs. Unexpectedly, the DNA-bound RNAP often restarted transcription, usually in reverse direction, thus producing an antisense transcript. Furthermore, we report evidence of this “secondary initiation” in live cells, using genome-wide RNA sequencing. These findings reveal an alternative transcription cycle that allows RNAP to reinitiate without dissociating from DNA, which is likely to have important implications for gene regulation.

## INTRODUCTION

In all organisms, gene transcription is usually viewed as initiating with the binding, assisted by accessory transcription factor proteins (Browning and Busby, 2004; Sikorski and Buratowski, 2009), of a RNA polymerase (RNAP) molecule from solution to a transcription promoter. In bacteria, the core RNAP first associates with an initiation factor, a sigma subunit, which confers the ability to recognize promoter DNA and initiate RNA synthesis (Gross et al., 1998). In the canonical bacterial transcription cycle, transcript synthesis concludes when release of the nascent RNA molecule from the polymerase is triggered by specific DNA sequences (intrinsic terminators) or by termination factors (e.g., the *E. coli* Rho protein) (Peters et al., 2011). While some studies suggest that RNAP dissociates rapidly from DNA upon intrinsic transcription termination, others suggest that a long-lived RNAP-DNA complex can persist after termination (Arndt and Chamberlin, 1988; Bellecourt et al., 2019; Larson et al., 2008; Yin et al., 1999).

Antisense transcription, which produces RNAs that have sequences at least in part complementary to ordinary “sense” gene transcripts, has been observed in organisms from bacteria to humans and is typically initiated from locations throughout the entire genome (David et al., 2006; Georg and Hess, 2018; He et al., 2008; Irnov et al., 2010). While the global biological significance of this pervasive antisense transcription has been questioned (Lloréns-Rico et al., 2016; Raghavan et al., 2012), antisense RNA production has demonstrated roles in regulating expression of many individual sense genes (Callen et al., 2004; Donovan et al., 2018; Georg and Hess, 2011; Lenstra et al., 2015; Sedlyarova et al., 2016; Shearwin et al., 2005; Tudek et al., 2015; Waters and Storz, 2009). The origins of antisense transcripts are incompletely understood. However, the relevant genetic elements, molecular mechanisms, and regulatory machinery are being explored (e.g., Brophy and Voigt, 2016; Murray et al., 2012; Sedlyarova et al., 2017).

In this study, we used single-molecule fluorescence microscopy *in vitro* to observe transcript production by bacterial RNAP and, significantly, also to follow the fate of the RNAP molecule after intrinsic termination of transcription. Under physiological ionic conditions, RNAP most often did *not* follow the canonical transcription cycle in which each recruitment event of a polymerase molecule to DNA can produce at most only one molecule of RNA primary transcript. Instead, the experiments reveal a frequently occurring alternative transcription cycle through which, as a consequence of its recruitment to a promoter, a single RNA polymerase molecule can produce multiple transcripts, including transcripts that are antisense to the first RNA molecule produced. In addition, we show evidence from end-enhanced genome-wide RNA sequencing suggesting that the alternative cycle is a widespread mechanism for synthesis of antisense transcripts in bacteria.

## RESULTS

### Observing single RNAP molecules from transcript initiation until after termination

To examine the behavior of individual molecules of RNA polymerase after transcription termination, we used a previously developed single-molecule fluorescence technique to study single-round transcription. In brief, we tethered fluorescent DNA molecules (DNA^488^) containing a promoter sequence, a 2.1 kbp transcription unit, and two consecutive intrinsic terminators to the surface of a glass flow chamber (Figure 1A). We incubated the surface with a solution containing σ^70^ holoenzyme made with core RNAP fluorescently labeled with a BG-549 dye on a SNAP tag on the carboxyl-terminal end of the beta subunit (RNAP^549^). Following open complex formation (Figure 1B, left), we initiated transcription by introducing 0.5 mM each ATP, CTP, GTP, and UTP at time *t* = 0, along with a Cy5-labeled oligonucleotide probe. The probe detects nascent transcript by hybridization to a repeat target sequence near the 5’ end of the RNA (Figure 1A). Analogously to previous experiments with labeled σ subunits (Friedman and Gelles, 2012; Harden et al., 2016), we observed the appearance of probe fluorescence spots that colocalized with RNAP^549^ and DNA^488^ spots, reflecting the hybridization of probe with the nascent RNA in individual transcription elongation complexes (ECs; e.g., *t* = 62 s in Figure 1B and Figure 1C, left). The probe spot typically later disappeared (e.g., *t* = 138 s in Figure 1B and 1C, left); this disappearance was scored as transcription termination since RNA release at intrinsic terminators is rapid (Yin et al., 1999) and the lifetime of transcript probe is not significantly reduced by photobleaching under these conditions (Figure S1A).

**Figure 1.**
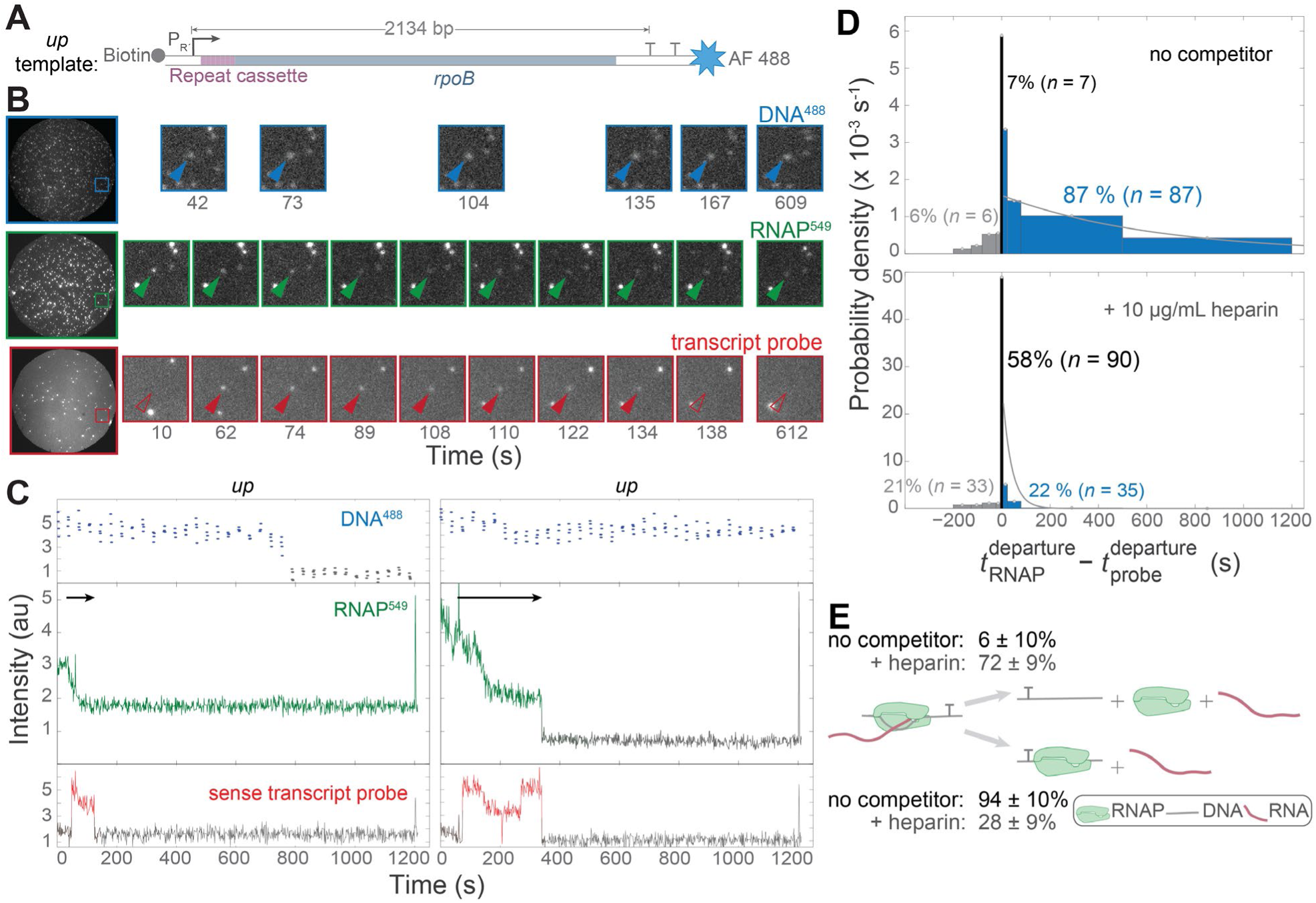
RNAP usually remains bound to the DNA template following transcript release at an intrinsic terminator. (**A**) *Up* transcription template. The template contains a wild type λ P_R’_ promoter region (blue, bent arrow) followed by seven tandem repeats of a 21 bp cassette (maroon), a partial sequence of *E. coli rpoB* coding region (gray) and two consecutive intrinsic terminators (T): λ T_R’_ and T7 T_E_, which have termination efficiencies *in* vitro of 93-95% and 88 ± 2%, respectively (DeVito and Das, 1994; Rees et al., 1997; Reynolds et al., 1992). Biotin is positioned upstream (“*up*”) of promoter so that RNAP moves away from the streptavidin-coated slide during transcription. (**B**) Left: Images (65 × 65 µm) of the same microscope field of view of DNA^488^ (blue), RNAP^549^ (green) and Cy5-transcript hybridization probe (red). Right: magnified views of the marked regions at various times during the experiment; NTPs were introduced at time *t* ∼ −10 s. Blue arrows mark the location of a DNA spot, green and red arrows mark the surface location in the other images, with presence (filled arrows) and absence (open arrows) of a co-localized fluorescence spot indicated. (**C**) Example fluorescence emission records from the locations of two DNA spots from the same experiment. Gray color marks intervals during which no fluorescent spot was seen. Arrows mark intervals of transcript elongation. Left: RNAP remains after probe departs (data from marked molecule in (B)) Right: RNAP and probe depart simultaneously. (**D**) Normalized histogram of RNAP^549^ departure time relative to Cy5-transcript probe departure from the same DNA spot for elongation reactions in the absence and presence of heparin. RNAP^549^ spot departed either before (gray), within 4 s of (black), or after (blue) transcript probe spot departure. The 4 s threshold was chosen because it is the maximum interval between consecutive frames. Gray curves are single exponential fits to the RNAP departure times following probe departure (see *Methods*). (**E**) Reaction scheme indicating the fraction of terminating ECs for which RNAP^549^ retains association with DNA after termination, calculated from the data in (D). See also Figure S1

### RNAP almost always remains bound to DNA after termination

In two replicate experiments, we observed a total of 100 molecules in which core RNAP^549^ fluorescence was visible when the probe spot appeared. Of these, 94% retained RNAP^549^ spots at the time of termination indicated by transcript release. Consistent with our earlier study with labeled σ^70^, most (87/100) RNAP^549^ molecules did not dissociate upon termination (Figure 1D, top). Instead, most persisted after transcript departure and eventually dissociated with a mean lifetime of 1140 ± 240 s (after accounting for photobleaching; see *Methods* and Figure S1B). Thus, nearly all RNAP that terminates under the conditions of these experiments stays associated with DNA after transcript release at an intrinsic terminator (Figure 1E). We previously showed that σ^70^-containing ECs behave similarly: on this same template 21% of ECs reached the terminator with bound σ^70^ and in the majority (74%) of these σ^70^ remained associated with DNA after termination (Harden et al., 2016). Taken together these data imply that both RNAP and σ^70^RNAP persist on DNA after transcription termination, usually for hundreds of seconds.

The presence of long-lived, DNA-RNAP complexes after termination is surprising. Heparin is a polyanion that can disrupt early promoter DNA-RNAP complexes in the initiation pathway (Ruff et al., 2015). When we added 10 µg/mL heparin together with the NTPs, we still observed transcript production from the open complexes as expected, but now most RNAP molecules dissociated from DNA within 4 s of transcript departure (Figure 1D, bottom). Those that did persist showed a characteristic lifetime (38 ± 10 s) greatly reduced relative to that in the absence of heparin. The observation that a polyanion competitor can dramatically reduce retention of RNAP on DNA after termination suggests that the retained RNAP interacts primarily with the DNA backbone without the more extensive contacts with DNA bases that occur in open complexes and ECs.

### RNAP can diffuse along DNA after termination

In our single-molecule experiments, transcript release was almost invariably preceded by a gradual decrease in RNAP^549^ fluorescence intensity (Figure 1C, black arrows). This decrease is expected, because during transcript elongation RNAP moves along the DNA so that its time-averaged distance from the chamber surface increases, decreasing the intensity of TIRF excitation and leading to reduced emission. A systematic increase in spot width was also observed, consistent with the idea that the intensity changes are due to net translocation of the elongation complex along DNA and result in increased Brownian motion of the DNA-tethered RNAP^549^ (Figure S1C, D, and E) (May et al., 2014). In an experiment with inverted DNA (i.e., with the biotin tag placed at the downstream end of the DNA), we instead observed increasing RNAP^549^ fluorescence during transcript probe co-localization, as predicted (Figure S1F).

In contrast to the gradual decrease in RNAP^549^ fluorescence observed before transcript release, we saw a different behavior after termination. After transcript release, we often (in roughly half of the Figure 1D blue population) saw episodes of rapid, bidirectional fluctuation in RNAP^549^ intensity (Figures 2A and S2A-I, teal). No correlated fluctuation was seen in DNA template fluorescence (e.g., Figure 2A, top), suggesting that the RNAP^549^ intensity fluctuations resulted from RNAP^549^ movements relative to the template DNA and not from transient sticking of the DNA to the surface. Similar large intensity fluctuations were not observed before or during the transcript probe signal, indicating that the movements are specific to the post-termination state. Measurements of RNAP^549^ position along the DNA derived from fitting the elongation portion of the record (Figure S2J) revealed that the post-termination fluctuations had the characteristics of a bounded one-dimensional random walk (Fig 2B, teal), frequently extending over the full ∼2 kbp template DNA length (e.g., Figure S2K). In some instances, the intervals of random motion were interspersed with periods of no apparent motion, during which the diffusion coefficient was zero within experimental uncertainty (e.g., Figure 2A, B purple).

**Figure 2.**
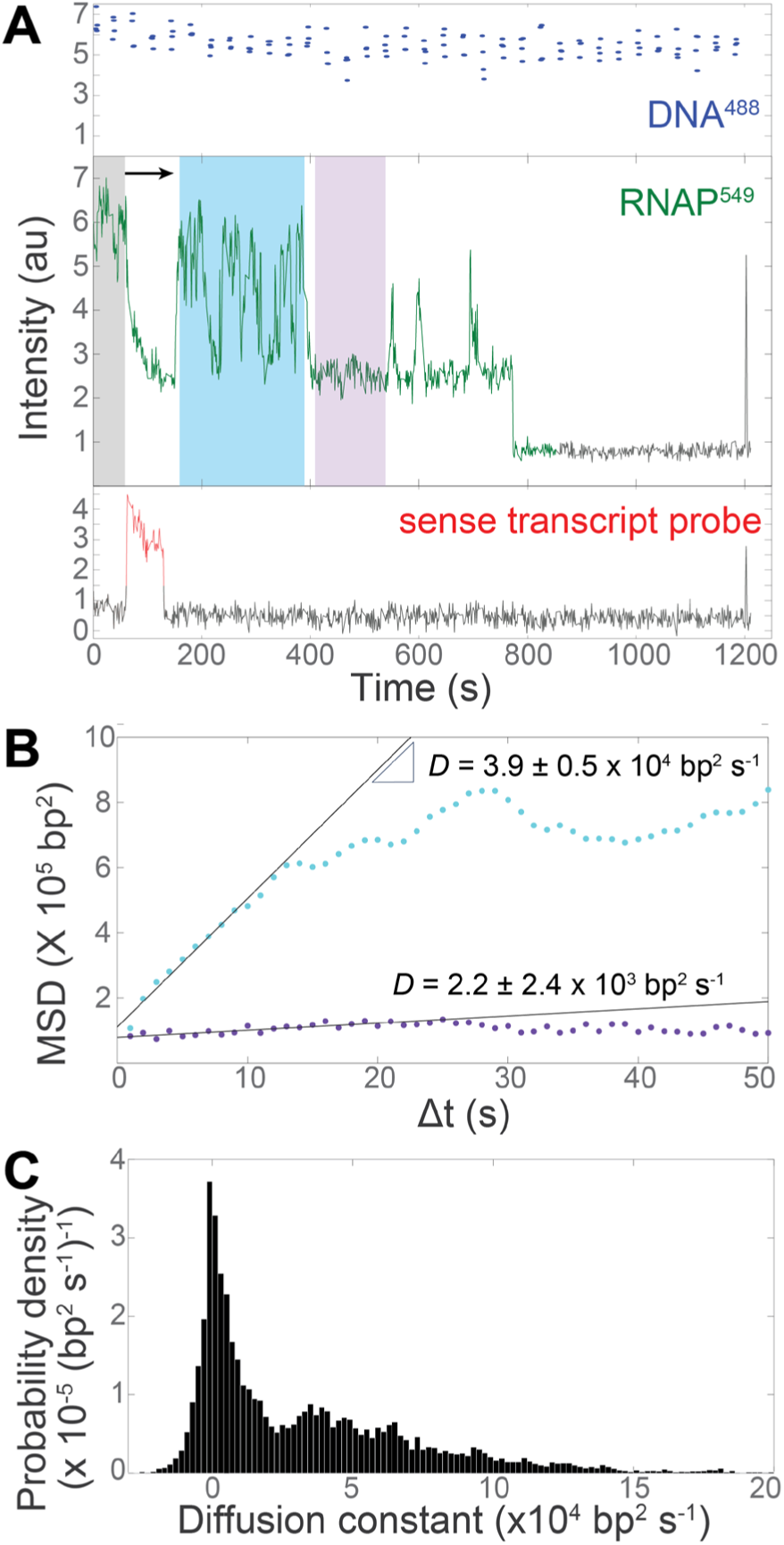
RNAP can diffuse along DNA after termination. (**A**) A single-molecule emission record, as in Figure 1C, for a different DNA spot location. Gray, teal, and purple highlight time intervals of high RNAP^549^ fluorescence before detection of transcript probe and post-termination intervals of fluctuating and low fluorescence, respectively. (**B**) Mean squared displacement (MSD) of RNAP^549^ position on DNA during the teal and purple intervals in (A). Linear fits to the first ten points of each MSD curve yield the effective diffusion coefficients over 10 s intervals, *D.* (**C**) Normalized histogram of *D* values measured separately for every 50 s window in *n* = 41 recordings of RNAP^549^ retained on DNA after termination (13,522 windows total). See also Figures S2 and S3.

Consistent with post-termination RNAP^549^ molecules switching between a state in which they slide randomly along DNA and a state in which they remain stuck at a fixed position, the distribution of measured diffusion coefficients was bimodal with peaks at ∼0 and ∼3.5 × 10^4^ bp^2^ s^-1^ (Figure 2C). Sliding diffusion coefficients of the latter magnitude are below the calculated upper limit for a protein of this size to randomly slide along the DNA helix (Blainey et al., 2009; Friedman et al., 2013). In a supplementary experiment in which a promoter-ablated mutant template was exposed to core RNAP^549^ in the absence of NTPs and σ^70^ (i.e., conditions in which neither promoter complexes nor elongation complexes should occur), qualitatively similar behavior was observed (Figure S3A and C), indicating that sliding/sticking motion on DNA may be an intrinsic property of core RNAP. In contrast, σ^70^RNAP holoenzyme exhibited typically much shorter interactions with DNA under the same conditions and showed few intensity fluctuations indicative of sliding (Fig S3 B and C). Since long duration sliding/sticking is seen with core RNAP and not with σ^70^RNAP, it is likely distinct from any sliding on DNA that might accompany promoter search by holoenzyme (Friedman et al., 2013; Wang et al., 2013).

### Post-termination RNAP-DNA complex can re-initiate transcription in the antisense direction

To test whether the sliding RNAP could rebind σ^70^ and initiate a new cycle of transcription, we performed further experiments, in which we introduced σ^70^ free in solution at the time of NTP introduction. The presence of free σ^70^ caused the behavior of most RNAP^549^ molecules retained after termination to change dramatically. Instead of the episodes of fast bi-directional sliding observed in the absence of σ^70^, we often observed slower unidirectional motion of RNAP^549^ in the opposite direction from the initial motion of transcript elongation: RNAP started near the promoter-distal end of the template DNA and moved towards the promoter (Figure 3A top left, gray arrow). Similar reverse unidirectional motion was also observed in a minority of cases even in the absence of free σ^70^ (e.g., Figure 3A, top right, gray arrow). In these cases, the reverse motion most often occurred after a brief period of sliding (e.g., teal regions in Figure 3A and Figure S2B), but it sometimes followed the forward motion with no discernable intervening sliding.

**Figure 3.**
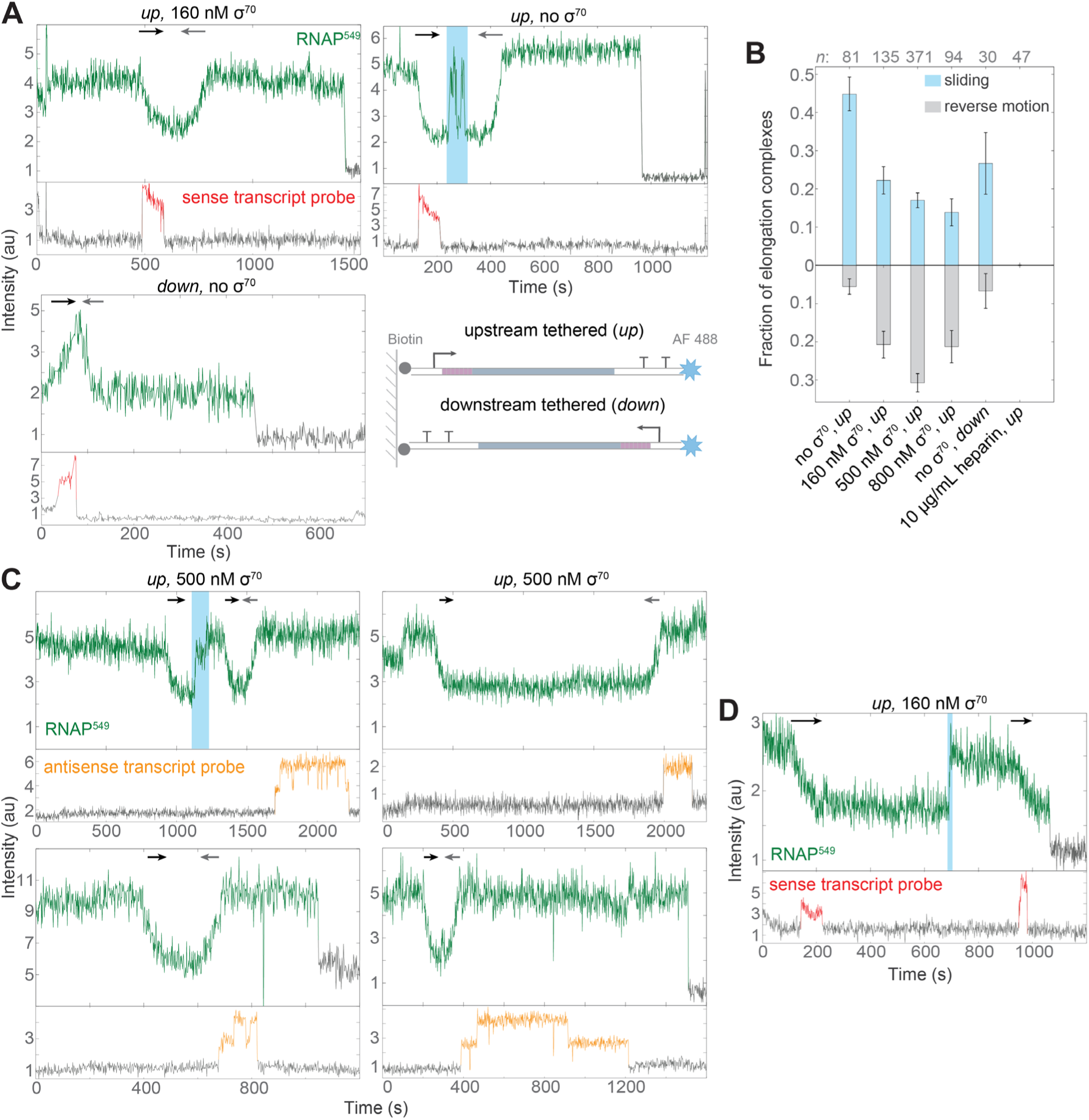
The post-termination RNAP-DNA complex can re-initiate transcription in the antisense direction. (**A**) Single-molecule emission records, as in Figure 1C and 2A, for three different DNA spot locations from three separate experiments: one with free σ^70^ in solution and *up* template (left), one with no free σ^70^ in solution and *up* template DNA (middle), and one with no free σ^70^ in solution and *down* template DNA (right). Black and gray arrows designate episodes of forward and reverse unidirectional motions corresponding to the directions of sense and antisense transcription, respectively. Teal indicates an interval of RNAP^549^ random sliding. Inset: Schematics of *up* and *down* templates using the same color scheme as Figure 1A. (**B**) Fraction (±SE) of RNAP^549^ molecules retained at termination (Figure 1D, black plus blue) that slide (teal) or exhibit reverse motion (gray) following termination under different experimental conditions with *up* or *down* templates. (**C**) Single-molecule emission records as in (A) with *up* template DNA and 500 nM free σ^70^ in solution and containing a transcript probe that is the reverse complement of that used in (A). (**D**) Single-molecule emission record as in (A) showing post-termination sliding (teal) followed by secondary initiation in the sense direction. See also Figure S4.

We performed analogous experiments using an otherwise identical template DNA molecule that was tethered to the surface by its downstream instead of its upstream end (Figure 3A, bottom). Forward followed by reverse motion was seen in the inverted template (e.g., Figure 3A, bottom right) in the expected directions (i.e., movement toward the surface followed by movement away from the surface). These observations show that reversal of direction is not restricted to the vicinity of the untethered end of the DNA.

The intensity changes corresponding to the forward (black arrows) and reverse (gray arrows) motions on an individual DNA usually exhibited mirror image shapes and similar durations (Figure 3A; Figure S4A). We hypothesize that the reverse motion reflects synthesis of an antisense transcript. Since core RNAP concentrations in solution are negligible in these experiments (Harden et al., 2016), and since we do not observe RNAP dissociation/re-association from DNA, antisense synthesis must be by the same RNA polymerase molecule that had just synthesized and terminated a sense transcript during the forward motion. This hypothesis predicts that a second initiation is required to produce the antisense transcript. Consistent with this prediction, the reverse motions were more frequent, occurring in up to 30% of elongation complexes, when σ^70^ (which is required for initiation) was present free in solution (Figure 3B). It should be noted that while the template lacks a known promoter for synthesis of an antisense transcript, it does contain an AT-rich sequence at the terminator that might act as a weak σ^70^ promoter. Both sliding and the reverse motions were absent when the polyanion heparin was present. Polyanions can disrupt the stable complexes that form between core RNAP and fully duplex DNA (Hinkle and Chamberlin, 1972; Melancon et al., 1983). Thus the heparin sensitivity suggests that the RNAP-DNA complex passes through a fully duplex-DNA intermediate (i.e., one with no open transcription bubble) prior to anti-sense initiation.

To check that the antisense transcript was made by the same RNAP molecule that just completed the sense transcript, we took advantage of the fact that even in highly purified *E. coli* RNAP preparations, each individual enzyme molecule has its own characteristic average transcript elongation rate (Adelman et al., 2002; Neuman et al., 2003; Tolić-Nørrelykke et al., 2004). Accordingly, we observed a broad range of characteristic intensity change rates for both sense and antisense transcription events (Figure S4B). However, the rate for a sense transcription event and the subsequent antisense event on the same DNA molecule were usually identical within experimental error, strongly suggesting that both were performed by the same individual RNAP molecule.

As an additional test of the idea that reverse motion is due to antisense transcription, we performed additional single-molecule transcription experiments (e.g., Figure 3C) using an antisense transcript probe complementary to the sense transcript probe used in Figures 1, 2, 3A and B. We found that 68% (78/114) of observed RNAP^549^ unidirectional motions towards the promoter were followed by subsequent antisense transcript probe co-localization, compared to just 2% (5/257) co-localization when no unidirectional motion towards the promoter was observed. These observations confirm that reverse motion was due to antisense transcript synthesis.

In the single-molecule experiments we sometimes observed sliding over long distances prior to secondary initiation (Figure 3A, top right). It is reasonable to ask whether RNAP ever slid back to the end of the DNA with the sense P_R’_ promoter and performed secondary initiation of a *sense* transcript. In rare cases (<2% of retained RNAP^549^ molecules), in the presence of free σ^70^ in solution, we observed retained RNAP^549^ molecules with intensity records indicating a brief period of sliding followed by re-initiation of transcription in the sense direction (Figure 3D, S4C), suggesting that secondary initiation can occur in either sense or antisense directions relative to primary initiation. Although antisense secondary initiation was preferred over sense in our data, that might be a characteristic of the DNA sequences used rather than an inherent feature of secondary initiation. Thus, free σ^70^ may confer onto DNA-bound core RNAP the capacity to locate and isomerize with promoter sequences after sliding hundreds of basepairs.

Antisense transcript was also detected in bulk transcription experiments at 500 nM free σ^70^ by RT-qPCR (Figures 4 and S5). Consistent with the single-molecule results indicating that the antisense transcript is made by polymerase molecules that have just completed a round of sense transcription, the bulk experiments showed that ablation of the promoter for the sense transcript reduced the concentrations of *both* sense and antisense transcripts (Figure 4). In this experiment, the ratio of antisense to sense transcript, 11 ± 2 %, was somewhat lower than seen in the single-molecule experiments (31 ± 2 % from 500 nM σ^70^ data in Figure 3B). This difference might result from transcriptional interference (Shearwin et al., 2005) caused by the multiple rounds of initiation possible in the RT-qPCR experiment; the design of the single-molecule experiment allowed only a single round. The RT-qPCR results were obtained with wild type RNA polymerase, confirming that the antisense transcript is not an artifact of the SNAP tagged and dye-labeled polymerase construct used in the single-molecule experiments. Taken together, the single molecule and bulk experiments show that antisense transcript synthesis on this template *in vitro* results from “secondary initiation” by RNAP molecules that first associated with the DNA through prior initiation at the sense promoter.

**Figure 4.**
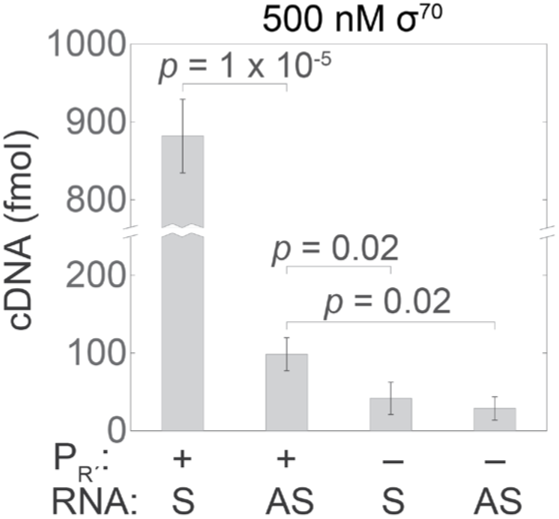
Amounts of both sense and antisense transcripts depend on the sense promoter. Measurements of the amounts of RNAs produced from bulk transcription reactions on templates with an unmodified (Figure 1A) or a scrambled (*Methods*) λ P_R’_ promoter. RNAs were reverse transcribed using sense (S) or antisense (AS) specific primers (Figure S5A) and the amounts of cDNA produced were measured by qPCR. Graph shows mean ± SE of three experiments. RT-qPCR controls are shown in Figure S5B. See also Figure S5.

### Secondary initiation of antisense transcripts *in vivo*

The forgoing experiments demonstrate that secondary initiation occurs *in vitro* with purified RNA polymerase and on a particular template DNA sequence but leave open the question of whether this phenomenon also occurs in living cells and on other template sequences. To investigate this, we used data from end-enriched RNA sequencing experiments that map the genomic positions of RNA 5’ and 3’ ends (Rend-seq; Lalanne et al., 2018). The secondary initiation hypothesis suggests that intrinsic terminators will be associated with nearby antisense initiation (Figure 5A). In *E. coli*, active intrinsic termination sites confirmed by Rend-seq data (Lalanne et al., 2018) were sometimes (in ∼20% of cases) accompanied by nearby (within 500 bp) RNA 5’ ends on the opposite strand indicating highly significant (> 12 standard deviations above the mean; see Methods) antisense initiation (Figure 5B, C, and D). These latter peaks were significantly more frequent near terminators than farther from them, indicating that the positions of sense terminators and antisense initiation were correlated (Figure S6A). Similar proximity was seen in data from *B. subtilis* (Figure S6B, C, and D), showing that it is a feature common to datasets collected from divergent species. Furthermore, the height of the antisense initiation peak was often increased in data from a *B. subtilis* strain with deletion of the Rho termination factor gene, consistent with prior observations that steady-state levels of antisense transcripts are greatly increased by Rho mutation or inhibition (Bidnenko et al., 2017; Lalanne et al., 2018; Nicolas et al., 2012; Peters et al., 2012). Both species display a consensus -10 box at sites of antisense initiation that is very similar to the -10 box at sense initiation sites (Shultzaberger et al., 2007), confirming that antisense initiation occurs at promoter-like sequences (Figure 5E, S6E; compare Dornenburg et al., 2010). In contrast, the antisense initiation peaks have -35 boxes different from those of sense initiation peaks when analyzed with the same algorithm (Bi and Rogan, 2006; see Methods). The -35 box is usually the most important sequence determinant in initial recruitment of RNAP to the promoter. The different sense and antisense -35 box sequences reported here may indicate that different sequences are optimal for different recruitment processes (e.g., binding of RNAP from solution for sense initiation vs. RNAP already bound nearby on DNA for antisense secondary initiation). Taken together, these RNA sequencing analyses show that antisense initiation occurs preferentially near some terminators, at discrete promoter-like sequences that differ from the sense promoter consensus. Such promoter-like sequences near terminators might be selected for (or, in other contexts, against) during genomic evolution. Sequencing does not follow individual RNAPs and thus cannot establish that sense and antisense RNAs are made sequentially by the same polymerase molecule. However, the data show antisense production that is consistent with the mechanism of secondary initiation (Figure 6) deduced from our experiments *in vitro.*

**Figure 5.**
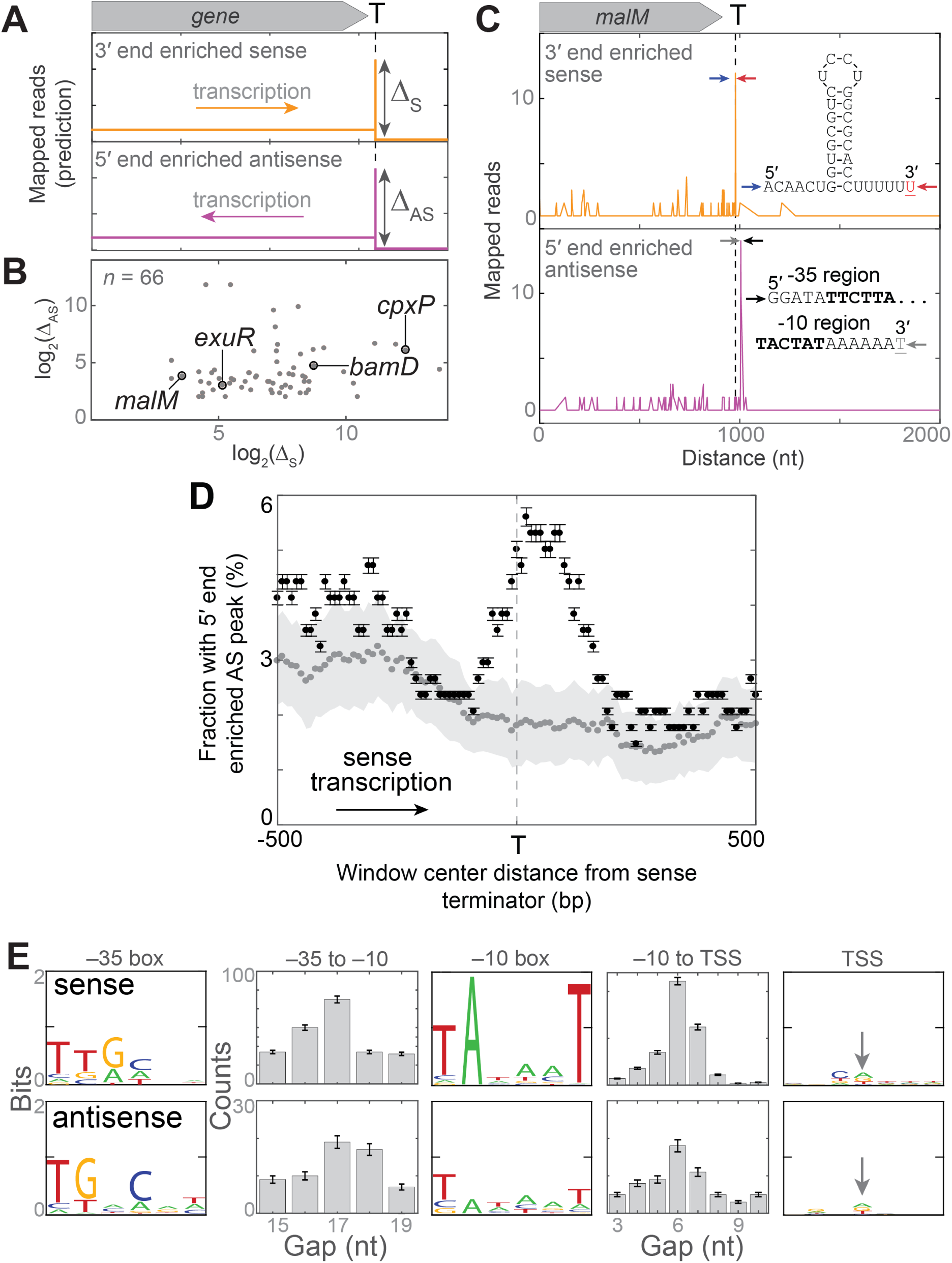
Secondary initiation of antisense transcripts *in vivo.* (**A**) Predicted Rend-seq signature of antisense secondary initiation at an intrinsic terminator. Idealized plot shows a genomic region near a terminator (T). Termination is indicated by a peak in 3′ end enriched sense reads (orange). The secondary initiation hypothesis predicts nearby 5′ end enriched antisense (magenta) reads, suggestive of antisense secondary initiation. Δ_*S*_ and Δ_*AS*_ are the relative amounts of sense termination and antisense initiation at particular genomic positions as estimated by the peak heights. (**B**) Peak heights from 66 (of 339 total) terminators detected in *E. coli* that show a substantial Δ_*AS*_ peak within 500 bp of the terminator Δ_*S*_ peak. Labels mark the genes shown in (C) and in Figure S6A. (**C**) Example of the phenomenon predicted in (A) observed in Rend-seq data from (Lalanne et al., 2018) near the terminator of the *E. coli malM* gene. Shown are the terminator RNA sequence with the peak of sense termination at the red nucleotide, and the promoter-like non-template strand DNA sequence with the peak of antisense initiation at the gray nucleotide. Arrows mark the positions of the displayed sequences in the Rend-seq data. (**D)** Antisense initiation peak frequency correlates with positions of sense terminators in the *E. coli* genome. Pooled data from 339 terminators between genes transcribed in the same direction (see *Methods*). Plot shows the fraction (± SE) of 200 nt-wide windows centered at the indicated distance upstream or downstream from the terminators that exhibit a peak of antisense initiation with *z*-score > 12 (black). Also shown is the mean ± SD of negative controls (gray) in which the same analysis was repeated 100 times each using 339 randomly selected locations in the *E. coli* genome that lack apparent terminators. These locations were restricted to those >700 nt from an annotated terminator and were on the sense strand of genomic regions containing at least three consecutive genes in the same orientation. In 100% of these 100 control replicates, the fraction at the terminator location with a 5’ end AS peak was < 3.9%, indicating that the difference between experimental data and controls was significant (*p* < 0.01). In this analysis, we used a smaller window size than in (B) to improve spatial resolution and a very stringent peak height criterion, *z* >12. This leads to detection of only the strongest peaks and shows that that these strong antisense peaks are preferentially found in a region ±200 nt from sense terminators. (**E**) Sequence consensus (illustrated as in (Shultzaberger et al., 2007)) for *n* = 250 strong sense promoters (top) and for the *n* = 66 terminator-proximal antisense initiation sites shown in (B) (bottom), all detected by Rend-seq; (Lalanne et al., 2018). Logos show the consensus sequences for the −35 box, −10 box, and transcription start site (TSS; arrow); histograms display the distributions of spacings between these elements. When the size of the - 10 box was expanded from six to 8, 9, or 10 bp, there was no strong evidence for extended -10 sequences. See also Figure S6.

**Figure 6.**
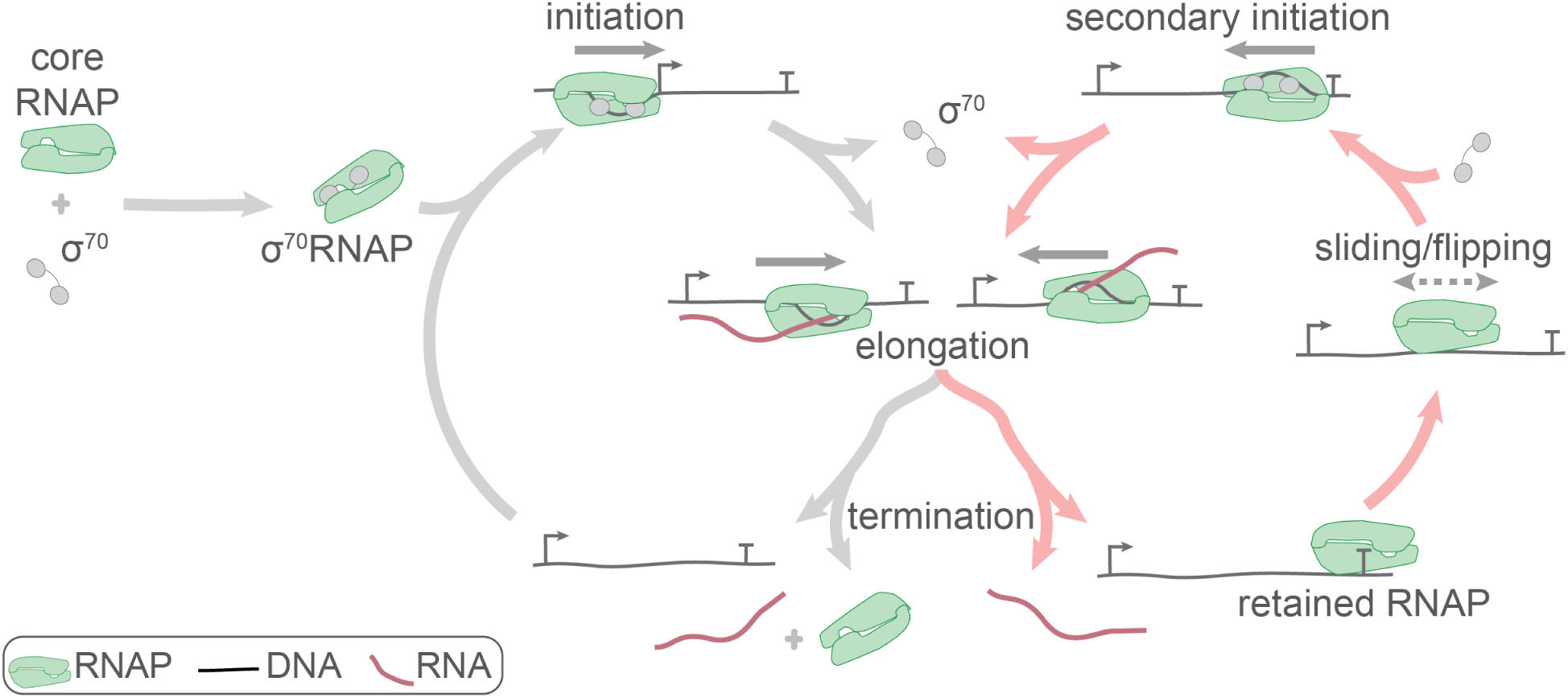
Proposed expanded bacterial transcription mechanism. RNAP retention after termination leads to an expanded pathway for transcript production, consisting of linked canonical (*gray*) and alternative (*red*) cycles. See text. Sequences that serve as antisense promoters and terminators are not shown.

## DISCUSSION

Taken together, our results suggest a new, expanded version of the bacterial transcription pathway (Figure 6), in which core RNAP retention on DNA after intrinsic termination can lead to synthesis of antisense (and possibly of additional sense) transcripts. In the canonical transcription cycle (Figure 6, gray arrows) holoenzyme formed by association of a sigma protein with core RNAP initiates at a sense transcription promoter and elongates a transcript. Sigma is released from most elongation complexes. The transcript and polymerase dissociate from template DNA upon reaching a sense transcript terminator (T) sequence. In the alternative cycle (Figure 6, red arrows), the termination process is different: core RNAP is retained on DNA after RNA is released at the terminator. This retained polymerase, which we assume is making only sequence non-specific interactions with the backbone of a fully base paired DNA, undergoes diffusional sliding along DNA. We further assume that the sliding RNAP molecule, like other sequence non-specific protein DNA complexes (see below), can occasionally flip its orientation on the DNA through transient dissociation and rapid rebinding. While in this sliding state, RNAP may bind a sigma factor, encounter a promoter-like sequence with orientation matching that of the polymerase, open a bubble in the DNA, and initiate a new transcript (secondary initiation) in a direction opposite to or the same as the direction of transcription before termination. Opposite-direction secondary initiation produces an antisense transcript. Each elongation complex is assumed to be capable of stochastically selecting either the canonical or the alternative cycle at the time of termination.

The nascent transcript dissociates rapidly (in ∼ 0.5 s) from RNAP when an EC reaches an intrinsic terminator (Yin et al., 1999). However, the rate of DNA release from RNAP at terminators is controversial; early studies produced indirect evidence for both rapid (seconds) and slow (minutes) release (Arndt and Chamberlin, 1988 and refs. cited therein). More recent studies show that at intrinsic terminators RNA release from RNAP occurs first and that an RNAP conformational change precedes subsequent DNA release (Bellecourt et al., 2019). In the single-molecule experiments, we directly measured the time between RNA release and DNA release and found that DNA release takes on average > 10 min in the absence of free σ^70^. This is consistent with earlier single-molecule observations (Yin et al., 1999) but superficially contradicts later work (Larson et al., 2008) which saw associations lasting only a fraction of a second. However, in those experiments the RNAP-DNA complexes were held under > 3 pN tension in an optical trap, a force predicted (see *Methods*) to move the sliding complexes we observe to the end of the DNA in 0.05 s. The optical trap data is consistent with our observations if one postulates that in those experiments the RNAP rapidly dissociated once it was pulled to the DNA end.

Early work characterizing RNAP-DNA interactions showed that core RNAP binds non-promoter DNA substantially more tightly than does σ^70^RNAP (Arndt and Chamberlin, 1988; Hinkle and Chamberlin, 1972; Melancon et al., 1983). However, there has been no known role for core RNAP-DNA interactions in the absence of the transcription bubble and nascent RNA present in an EC. Here, we show that a core RNAP-DNA complex is a transcription cycle intermediate that is often produced upon transcript release at one or both of the intrinsic terminators used in our experiments. These post-termination complexes are kinetically stable, and they often exhibit long-range sliding along DNA. Evidence that most or all of the sliding complexes contain core RNAP rather than holoenzyme includes: 1) the fraction of complexes that we see slide post-termination (Figure 3B) is much larger than the fraction that retain σ^70^ (∼21%; see Harden et al., 2016); 2) the observation that adding σ^70^ to the solution suppresses post-termination sliding (Figure 3B); and 3) that incubating core RNAP with promoterless DNA produces similar long-lived sliding complexes (Figure S3). The behaviors observed in the presence of free σ^70^ further suggest that after termination, the sliding core RNAP-DNA complexes can bind σ^70^ and re-initiate transcription. We speculate that this sliding-mediated secondary initiation represents a previously unknown biological function of the kinetically stable core RNAP-DNA interaction.

Our results suggest that after one round of transcription, RNAP can initiate a second round in the opposite direction without intervening dissociation and diffusion of the enzyme away from the DNA. This “flipping” presumably requires RNAP to rotate by 180 degrees about an axis normal to the DNA helix. Although flipping has not previously been reported for RNAPs, it has multiple precedents in other enzymes that slide on or move processively along nucleic acids (Ganji et al., 2016 and references cited therein; Comstock et al., 2015). In those enzymes, flipping is presumed to occur via undetectable brief dissociation limited to the microsecond/nanometer scale followed by rapid rebinding of the protein to the DNA (Ganji et al., 2016; Paramanathan et al., 2014). It is conceivable that in bacterial RNAPs, an α subunit C-terminal domain (Murakami, 2015) could increase the efficiency of flipping by flexibly tethering RNAP to DNA while it rotates.

Although secondary initiation has not previously been reported for bacterial RNAP, there is evidence that the same molecule of eukaryotic RNA polymerase III can re-initiate a second round of sense transcription at same promoter after termination of the first round (Dieci and Sentenac, 1996; Ferrari et al., 2004). While this re-initiation has been proposed to occur by a looping or “handing back” mechanism mediated by transcription factors, our results with bacterial RNAP suggest sliding as a possible alternative mechanism.

Antisense transcription is known to act through transcription interference and other processes to regulate specific genes in bacteria (Georg and Hess, 2011, 2018), and terminator/antisense promoter modules have been shown in synthetic genetic constructs to exert a general suppressive effect on transcript production from the upstream sense gene (Brophy and Voigt, 2016). Antisense transcription, including transcripts that initiate downstream of sense terminators, is pervasive in bacteria, but the mechanisms that give rise to it are not well understood and antisense promoter sequences are not well conserved (Raghavan et al., 2012). The retention after termination/sliding/flipping mechanism described here is noteworthy because antisense transcript production immediately follows and is coupled to the production of a sense transcript from the same gene by the same RNAP molecule. Thus, initiation at the sense promoter can directly produce an antisense transcript to down-regulate sense gene expression. This mechanism could provide a fast-acting negative feedback that suppresses spurious expression in bacteria without the time required for translation, serving a regulatory role similar to that reported in regulation of eukaryotic transcription (Lenstra et al., 2015). In addition, our observations raise the possibility that the presence of a sense promoter(s) near an intrinsic terminator could cause RNAP retained after intrinsic termination to do secondary initiation in the sense direction from the nearby promoters. This could serve as a gene coupling mechanism in which transcription from an operon could serve to activate adjacent promoters, leading to local regions of enhanced transcription in the bacterial genome. Further study will be required to elucidate the gene-specific roles of these molecular behaviors in living cells.

## Acknowledgments

We would like to thank members of the Landick, Block, DePace, Kondev and Gelles labs for insightful discussion as well as Clarissa Scholes for figure design. We are grateful to Larry Friedman and Johnson Chung for help with microscopy and data analysis. We thank Michael Rosbash (Brandeis) and Michael Carrel (Children’s Mercy Kansas City) for access to computational resources and software. This work was supported by grants from the G. Harold and Leila Y. Mathers Foundation, the National Institute of General Medical Sciences (R01GM081648, R35GM124732, R01GM044025, and R01GM38660), and the National Science Foundation (DMR-1610737 and MRSEC DMR-1420382), Pew Biomedical Scholars Program, Searle Scholars Program, Sloan Research Fellowship, Smith Family Awards, National Science and Engineering Research Council of Canada Fellowship, Howard Hughes Medical Institute International Student Research Fellowship;

## Author contributions

Harden, Chamberlain, Herlambang, Kondev, and Gelles designed the research; Harden, Chamberlain, Herlambang, and Lalanne performed the research; Lalanne, Wells, Li, and Hochschild contributed unpublished data or new reagents; Harden, Chamberlain, and Herlambang analyzed the data; Harden, Chamberlain, Kondev and Gelles drafted the manuscript and all authors contributed to writing the final version.

## METHODS

### Template DNA and oligonucleotides

Circular transcription templates were the plasmids pCDW114 (GenBank accession no. KT326913) and pCDW116. Plasmid pCDW116 has the same sequence as pCDW114 but the P_R’_ –35-box TATTGACT in pCDW114 was mutated to CAGGCGCT. Linear transcription templates (*up* and *down* DNA^488^, Figs 1A and 3A) were synthesized by PCR from plasmids pCDW114 using the primers described previously (Harden et al., 2016). The template lacking a P_R’_ promoter sequence (Figures 4, S3, and S5) was synthesized in the same way using plasmid pCDW116. The 20nt Cy-5-labeled sense and antisense transcription probes were 5’-GTG TGT GGT CTG TGG TGT CT/3Cy5Sp/-3’ and 5’-AGA CAC CAC AGA CCA CAC AC/3Cy5Sp/-3’, respectively (IDT, Coralville, IA).

### Proteins

*E coli* Core RNAP (αββ’ω) with a SNAP tag on the c-terminus of β’ (RNAP-SNAP) and wild type σ^70^ protein was expressed and purified as described (Tetone et al., 2017). RNAP-SNAP was labeled with the DY-549 dye, yielding RNAP^549^, as follows: 20 μL of 15 μM RNAP-SNAP was dialyzed into 3 L of labeling buffer (10 mM Tris-HCl, pH 8.0, 40 mM KCl, 5 mM MgCl_2_, 20 μM ZnCl_2_ and 1 mM dithiothreitol (DTT)) at 4 °C for 4 h. The resulting product (typically 50 – 100 μL of 5 – 20 μM of protein) was mixed with an equimolar amount of SNAP-Surface 549 (New England Biolabs; 1 mM in DMSO) and incubated at room temperature for 30 min, then mixed with an equal volume of labelling buffer supplemented with an 60% glycerol to yield RNAP^549^ in reconstitution buffer (10 mM Tris-HCl, pH 8.0, 30% glycerol, 0.1 mM EDTA, 100 mM NaCl, 20 mM KCl, 20μM ZnCl_2_, 3 mM MgCl_2_ and 0.6 mM DTT). The preparation was flash frozen in liquid N_2_ and stored at −80 °C.

σ^70^RNAP^549^ holoenzyme was prepared by incubating equimolar σ^70^ and RNAP^549^ in reconstitution buffer at 37°C for 10 min and then stored at −20 °C for up to 3 hr before use.

### Single molecule transcription experiments

Single-molecule total internal reflection fluorescence microscopy was performed at excitation wavelengths 488, 532 and 633 nm, for observation of DNA^488^ template, RNAP^549^ and Cy5-transcript probe, respectively, as described (Friedman and Gelles, 2012); focus was automatically maintained as described (Crawford et al., 2008). Transcription reactions were conducted as described (Harden et al., 2016). Briefly, single-molecule observations were performed in glass flow chambers (volume ∼20 µL) passivated with succinimidyl (NHS) polyethylene glycol (PEG) and NHS-PEG-biotin (Laysan Bio Inc.; Arab, AL) as described (Friedman and Gelles, 2012). Streptavidin (#21125; Life Technologies; Grand Island, NY) was introduced at 220 nM in wash buffer (50 mM Tris-OAc, 100 mM KOAc, 8 mM MgOAc, 27 mM NH_4_OAc, 0.1 mg mL^-1^ bovine serum albumin (BSA) (#126615 EMB Chemicals; La Jolla, CA), pH 8.0), incubated 45 s, and washed out (this and all subsequent wash out steps used two flushes each of four chamber volumes of wash buffer). The chamber was then incubated with 50 pM AF488-DNA in wash buffer for ∼2 min and washed out. Next, locations of surface-tethered AF488-DNA molecules were recorded by acquiring four 1 s images with 488 nm excitation at a power of 350 µW incident to the objective lens (Crawford et al., 2008).

For transcription reactions σ^70^RNAP^549^ holoenzyme was introduced into the chamber at 1 nM in transcription buffer (wash buffer supplemented with 3.5% w/v PEG 8,000 (#81268; Sigma-Aldrich; St. Louis, MO), 1 mg mL^-1^ BSA, and an O_2_-scavenging system (Friedman et al., 2006), incubated for ∼10 min, and washed out. Finally, we started image acquisition (iterations of thirty 1 s exposures to simultaneous 532 and 633 nm excitation, each at 200 µW, followed by four 1s exposures to 350 µW 488 nm excitation) and initiated transcription by introducing transcription buffer supplemented with 500 µM each of ATP, CTP, GTP and UTP, and 10 nM Cy5-probe.

Image analysis was done using custom software and algorithms for automatic spot detection, spatial drift correction and co-localization as described (Friedman and Gelles, 2015).

### Bulk transcription experiments

Open-promoter complexes were formed by combining 8.8 nM unlabeled σ^70^-SNAP-RNAP holoenzyme with 8 nM of DNA template in 50 µL of transcription buffer supplemented with 660 nM σ^70^ and incubated for 5 min. Transcription was then initiated by the introduction of 500 μM each of ATP, CTP, GTP and UTP. The reaction was allowed to proceed for 40 min at room temperature; at that time total RNA was purified using RNeasy mini Kit (Qiagen; Cat No. 74104) column and protocol including on-column RNase-free DNase digestion (Qiagen; Cat No. 79254) and eluted into 30 μL RNase free water.

### RT-qPCR

First strand complementary DNA (cDNA) was synthesized in a 25 µL reaction containing 12.5 µL sample RNA, 2 pmol strand-specific cDNA primer (Figure S5A) and 200 units SuperScript IV reverse transcriptase (ThermoFisher; Cat No.18090010) in RT buffer (50 mM Tris-HCl, pH 8.3, 50 mM KCl, 3 mM MgCl_2_, 10 mM DTT, and 1 mM each dATP, dCTP, dGTP, dTTP) and incubated according to the SuperScript IV reverse transcriptase protocol. cDNA product was diluted 1:2 into TE buffer (10 mM Tris-HCl pH 8.0, 0.1 mM EDTA). qPCR was conducted using qPCR primers chosen to amplify the cDNA (Figure S5A) in 20 μL reactions containing 4 µL diluted cDNA, 0.5 μM primers, 0.2 µL Herculase II Fusion DNA Polymerase (Agilent Technologies; Cat No 600675), and Sybr Green (ThermoFisher) at the manufacturer’s recommended concentration. cDNA synthesis reactions were performed on three different days; subsequent to each cDNA reaction qPCR was performed in triplicate on each sample. On each day, sense and antisense standard curves were measured from nine qPCR reactions containing known amounts of target sequence double stranded DNA (6 × (10^2^, 10^3^, 10^4^, 10^5^, 10^6^, 10^7^, 10^8^, 10^9^ or 10^10^)) molecules. Sense or antisense cDNA copy number in each qPCR reaction was calculated using parameters derived from fitting the corresponding standard curve. Mean qPCR amplification efficiency was 104 ±4%.

### Data analysis

#### Characteristic lifetime of RNAP^549^

To measure the characteristic lifetime of retained RNAP^549^, we jointly fit to an exponential probability distribution the measured lifetimes of retained RNAP^549^ that terminated by disappearance of the fluorescent spot and those that were censored by halting image acquisition using the maximum likelihood algorithm, yielding the reciprocal time constant *k*_*obs*_ (Harden et al., 2016). The dissociation rate of retained RNAP^549^, *k*_RNAP_, was computed by *k*_RNAP_ = *k*_obs_ – *k*_PB_ where *k*_PB_ is the rate of RNAP^549^ photobleaching (Figure S1B) and the characteristic RNAP lifetime was calculated as 1 / *k*_RNAP_. Errors were calculated by bootstrapping as described (Friedman and Gelles, 2015) and error propagation.

### Measurement of RNAP^549^ position on template

We used location-specific calibration curves at the position of each DNA molecule to convert measured RNAP^549^ fluorescence intensity to position along the DNA contour. To define the calibration curve, we first fit the RNAP^549^ fluorescence record during the period of steady state elongation (Fig S2J, black arrow) to the expression

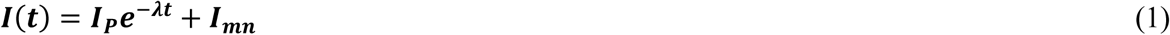

where *I*_*P*_ and *I*_*mn*_ are the fluorescence intensity of the promoter-bound RNAP^549^ and the mean magnitude of the background fluorescence as depicted in Figure S2J, and the fit parameter *λ* is the decay constant (Figure S2J, blue curve). We assumed the rate of elongation was constant (Adelman et al., 2002; Neuman et al., 2003; Tolić-Nørrelykke et al., 2004), yielding the relationship

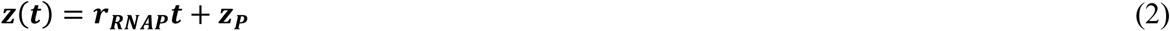

where *z*(*t*) is the position of the polymerase along the contour of the DNA during elongation, *r*_*RNAP*_ is the rate of RNAP elongation, and *z*_*p*_ is the position of the promoter along the DNA contour. Taking the time of probe release as the time of termination, we measured the fluorescence intensity at termination, *I*(*t*_*T*_) = *I*_*T*_ (Figure S2J), and used it to compute *r*_*RNAP*_ by combining Eqn. 1 and Eqn. 2 and using the known position of the terminator along the DNA contour *z*_*T*_:

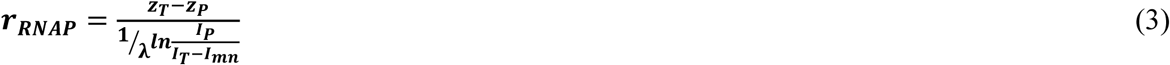

Finally, combining Eqns. 1 and 2 yields an expression relating the time-dependent position of the polymerase on the DNA contour to the measured time-dependent fluorescence intensity *I*(*t*) after termination

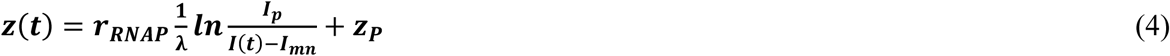

in terms of known and measured parameters. An example record of *z*(*t*) is shown in Figure S2K.

### Identifying sliding and antisense transcription behavior

To measure the fractions of retained RNAP^549^ molecules that exhibited post-termination sliding or antisense transcription (Figure 3B), we analyzed all RNAP^549^ fluorescence emission records that displayed sense transcript elongation as judged by fluorescence intensity changes. Sliding was scored if any 50 s time window following the RNAP^549^ elongation signature contained a measured diffusion coefficient, *D*, of 2.2 × 10^4^ bp^2^ s^-1^ or greater. This *D* value corresponds to the local minima of the saddle point in Figure 2C. Antisense transcription was scored if a region of the RNAP^549^ intensity record after sense transcript elongation completed exhibited a visible antisense elongation profile that when fit had an exponential time constant between 0.002 and 0.04 s^-1^.

### Estimate of post-termination RNAP drift velocity under force

Previous work by Larson et al. (2008), employed an optical trapping assay featuring bead-tethered RNAP undergoing steady-state elongation on surface-tethered template DNA, with tension (as small as 3 pN) imposed between the two by the trap. Upon RNA reaching the position of an intrinsic terminator on the DNA, dissociation of RNAP from DNA was detected as loss of the mechanical linkage between bead and surface. These data were interpreted as sub-second dissociation of RNAP from DNA after intrinsic termination. Here we claim that our observation of a long-lived (hundreds of seconds), DNA-bound sliding RNAP state following intrinsic termination is fully consistent with the sub-second dissociation seen under applied force.

The Einstein–Smoluchowski equation relates the one-dimensional diffusion constant of a particle, *D*, to the drift velocity, *v*_*d*_, under an external force, *F*:

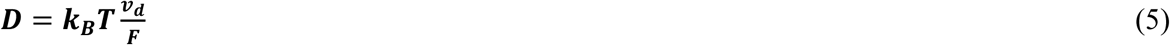

where *k*_*B*_ is Boltzmann’s constant and *T* is temperature. Solving Eqn. 5 for *v*_*d*_ and evaluating the expression using our measured diffusion constant of RNAP on DNA, *D* = 4 × 10^4^ bp^2^ s^−1^ = 4 × 10^3^ nm^2^ s^−1^, the minimum external force imposed on RNAP relative to DNA in (Larson et al., 2008), *F* = 3 pN, and *T* = 300 K, yields the drift velocity of the post-termination RNAP in the sliding state under the external force imposed by the optical trap: *ν*_*d*_ = 3 × 10^3^ nm s^−1^. At this drift velocity, RNAP in the optical trap assay will be pulled along the DNA from the position of the terminator to where it could slide off of the blunt end (∼150 nm) in ∼0.05 s, consistent with the rapid dissociation observed in those experiments.

### Genome-wide effects of intrinsic terminators on antisense transcript production

To establish a reference set of terminators for analysis, we used sets of 630 *E. coli* and 1,486 *B. subtilis* terminators with terminator function *in vivo* established by experimental data on wild-type strains (Lalanne et al., 2018). To ensure that the identification of 5’ ends was not affected by peak shadows near the ends of convergent genes (Lalanne et al., 2018), we restricted our analysis to a subset of *n* = 339 (*E. coli*) or 726 (*B. subtilis*) terminators for which the nearest upstream and downstream genes were annotated in the reference genome NC_000913.2 (*E. coli*) or NC_000964.3 (*B. subtilis*) to be in the same orientation as the terminator. To quantify sense termination, we first defined *k*_*max*_ as the peak number of 3’ end-enriched reads mapped to the same strand as the terminator in a 10 bp region around each terminator. The magnitude of sense termination was taken to be Δ_*S*_ = log_2_(*k*_*max*_) (Figures 5B and S6B). To estimate the effect of the terminators on antisense transcript production, we first defined *k*_*max2*_ as the peak count of 5’ end-enriched reads mapped to the opposite strand in a ±500 bp region around each terminator. The magnitude of antisense initiation was taken as Δ_*AS*_ = log_2_(*k*_*max2*_). Antisense initiation peaks were identified by *z*-score transformation as described (Lalanne et al., 2018); a threshold of *z*-score > 12 was used to select strong peaks (*n* = 66 *E. coli* or *n* = 117 *B. subtilis* Δ*rho* terminators met this criterion).

### Consensus promoter sequence determination

To determine consensus sequence of the TSS (Figures 5E and S6E), at each antisense initiation peak, we first measured the information content of the nucleic acid sequence (Shultzaberger et al., 2007) in a ±3 bp window centered on each peak. To determine the consensus sequence of the −10 box, as well as the distribution of gap lengths between the TSS and −10 box, we used BIPAD, a web server for modeling bipartite sequence elements with variable spacing (Bi and Rogan, 2006). After substituting 7 A nucleotides for positions +1 through +7 (relative to the TSS at +1), we fit positions −20 to +7 (BIPAD parameters: gap range, 3-10 bp; widths of sequence elements, 6 bp and 7 bp; 1000 runs). To determine the consensus sequence of the −35 box, as well as the distribution of gaps between the −10 box and −35 box, positions −43 to −3 were fit (BIPAD parameters: gap range, 15-19 bp; two sequence-element search; widths, 6 bp and 6 bp; 1000 runs). For comparison, the same analysis was used on sets of *coli* and *B. subtilis* sense initiation peaks detected in Rend-seq data. These sense initiation peaks were identified by peak *z*-score > 12 (Lalanne et al., 2018) in wild-type cells.

## SUPPLEMENTARY FIGURES

**Figure S1.**
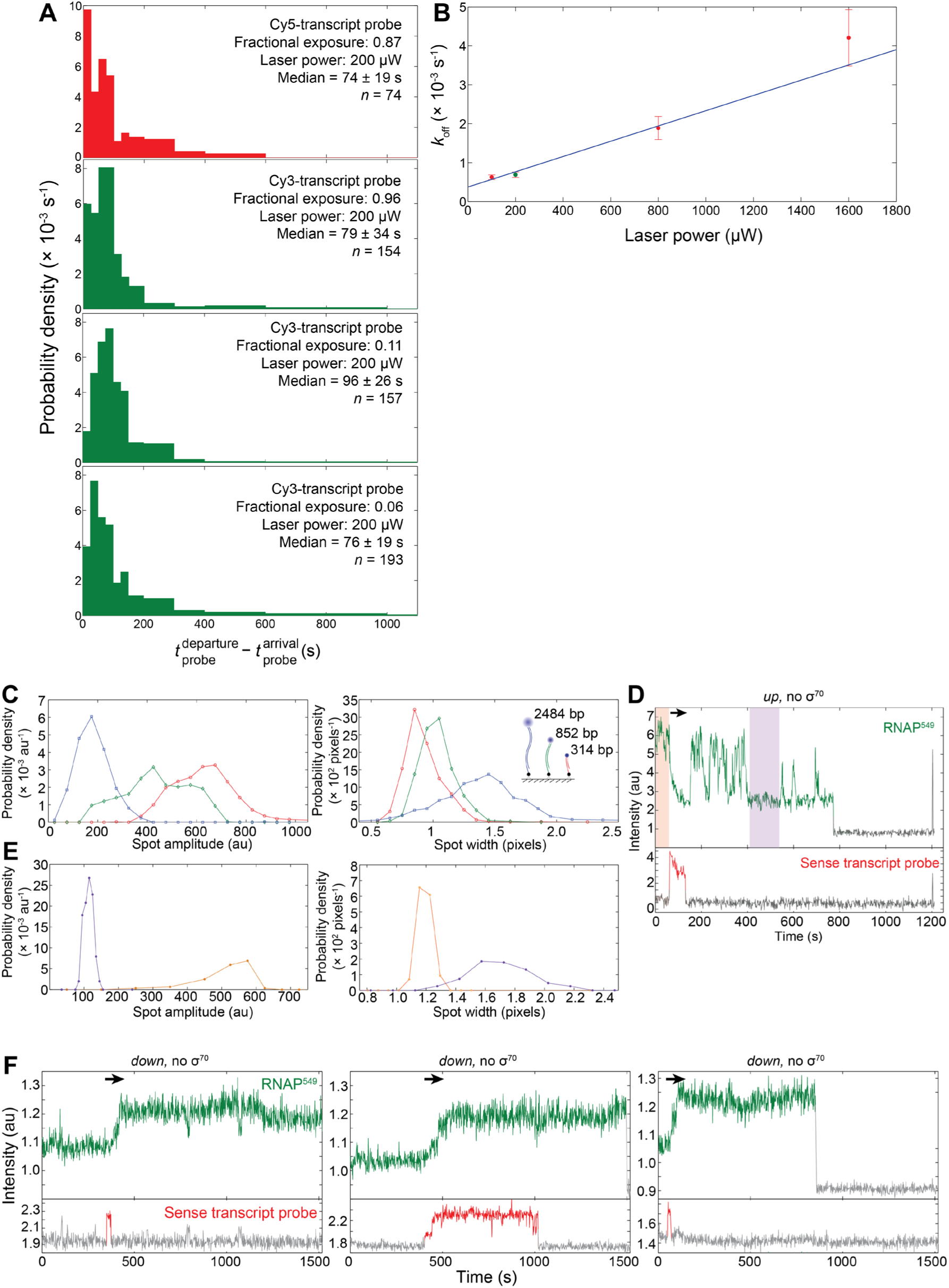
Validation of single-molecule experiments to observe pre- and post-termination RNAP-DNA complexes. Related to Figure 1. (**A, B**) Effects of laser exposure on single-molecule photobleaching lifetimes of fluorescently tagged species. (**A**) Normalized histograms of transcript probe lifetimes with either a Cy5 (red) or Cy3 (green) labels on the probe, measured at the indicated fractional exposures (i.e., the fraction of the time the sample is exposed to the excitation laser). A Kolmogorov-Smirnov test between each distribution (*p* = 0.95) failed to reject the null hypothesis that each of the distributions were selected from the same continuous distribution. The red histogram corresponds to the conditions used in this study; the green histograms are taken from (Harden et al., 2016). The similarity of the lifetime distributions over a 16-fold range of fractional exposure suggests our measurements of Cy5-probe lifetime are not significantly limited by photobleaching (Harden et al., 2016). (**B**) Measurements of the first-order fluorescent spot disappearance rate constant *k*_off_ of RNAP^549^ in open-promoter complexes with template DNA at varying excitation laser powers. The point at 200 µW (green) corresponds to the power used in the experiments reported in the study. Linear fit (line) yielded intercept (3.7 ± 0.4) × 10^-4^ s^-1^ (the open complex dissociation rate after photobleaching correction) and slope (2.0 ± 0.8) × 10^−6^ s^-1^ µW^-1^, corresponding to a photobleaching rate at 200 µW of *k*_PB_ = (7 ± 2) × 10^-4^ s^-1^ or photobleaching lifetime τ = 1300 ± 300 s, significantly more than the median lifetime of the transcript probe (75 ± 18 s). (**C, D, E**) Tethered fluorophore motion reports movement of RNAP along DNA. **(C)** Control experiment with three different lengths of dye-labeled DNA confirms a previous report (May et al., 2014) that florescence spot intensity and width change systematically with DNA tether length due to tethered fluorophore motion (TFM). Three DNA species of lengths 2484 bp (blue), 852 bp (green) and 314 bp (red), each modified with an AF488 dye at one end and biotin at the other, were sequentially introduced into a streptavidin-derivatized flow chamber. After each introduction, the locations of molecules of that length were recorded for subsequent analysis. A recording of a single field of view with all three molecular species present was then analyzed by fitting each fluorescent spot with a two-dimensional Gaussian (Friedman and Gelles, 2015). Histograms show the resulting spot amplitudes (left) and width (i.e., the standard deviation of the Gaussian; right) classified by DNA type. Analysis was conducted on 10 images (each 1s exposure time) containing 70 (blue), 53 (green) and 102 (red) DNA molecules, producing 700, 530, and 1020 measurements, respectively in the three distributions. (**D**) The same experimental record shown in Figure 2A. Shading indicates two time intervals chosen to be prior to transcript elongation (salmon) and after termination (purple) as judged by the transcript probe signal. (**E**) Histograms of spot amplitude (left) and width (right) demonstrating detection of RNAP^549^ movement on DNA during the single-molecule transcription event depicted in (D). Spot images used to produce the salmon (*n* = 61) and purple (*n* = 101) histograms were drawn from the indicated time intervals in (D). Before elongation, RNAP^549^ is located closer to the surface than it is during the selected interval after termination, as demonstrated by the lower amplitude and greater width in the purple distributions relative to the salmon. (**F**) Example records illustrating RNAP transcribing a template DNA that is inverted relative to the template orientation used in Figure 1. Records selected from an experiment with *down* (*Methods*) template DNA showing RNAP^549^ and Cy5-transcript probe emission co-localized with 3 different DNA spots, plotted as in Figure 1C. Gray color marks intervals during which no fluorescent spot was detected. Arrows mark intervals of transcript elongation. This template is attached to the surface in the opposite orientation from that in Figure 1A, and the fluorescence intensity increases, rather than decreases, during transcript elongation as expected.

**Figure S2.**
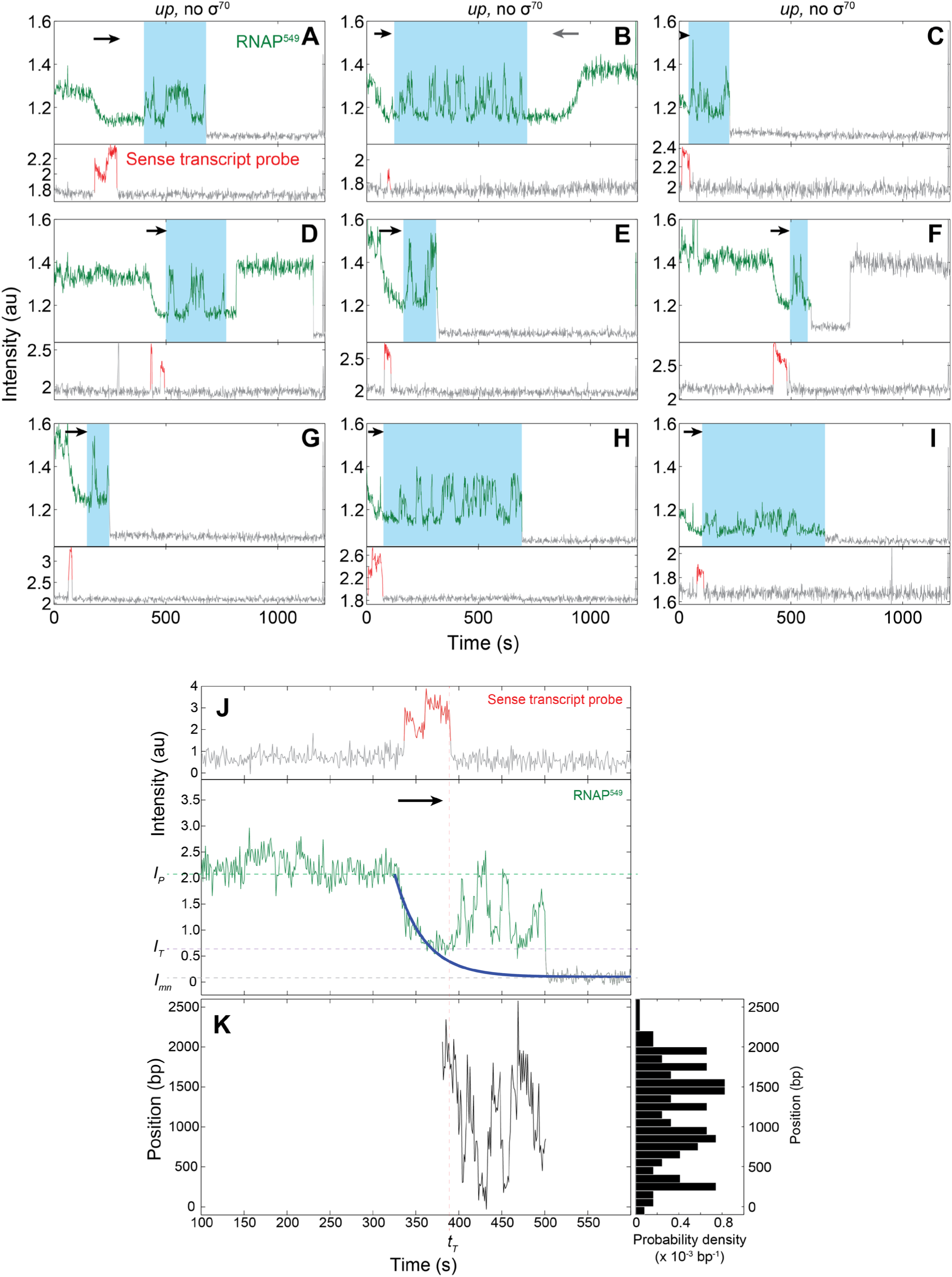
Example records illustrating RNAP sliding on DNA after termination and measuring retained RNAP^549^ position on template DNA. Related to Figure 2. (**A-I**) Records selected from the experiment in Figures 1 and 2 showing RNAP^549^ and Cy5-transcript probe emission co-localized with nine different DNA spots. Gray color marks intervals during which no fluorescent spot was seen. Black and gray arrows designate episodes of forward and reverse unidirectional motions corresponding to the directions of sense and antisense transcription, respectively. Teal indicates intervals of RNAP^549^ random sliding. (**J**) Computing the position of retained RNAP^549^ on template DNA. Single-molecule emission records, as in Figures 1, 2, 3 and S1. The time interval in the RNAP^549^ record indicated by the black arrow is fit to a single exponential decay model (solid blue curve, *Methods*). The dashed lines mark the time of termination *t*_T_ (red), the RNAP^549^ fluorescence intensity while bound to promoter *I*_P_ (green) and at the time of termination *I*_T_ (purple), and the mean background fluorescence when no RNAP^549^ spot is present *I*_mn_ (gray). Right, schematic of a model of RNAP^549^ position on the template DNA during constant-velocity elongation (*Methods*) accompanied by a depiction of template DNA indicating the approximate locations of the promoter (*z*_P_) and terminator (*z*_T_) regions. (**K**) Position of RNAP^549^ after termination calculated using the calibration depicted in (J) (*Methods*). Right, normalized histogram of RNAP^549^ position during the interval plotted at left.

**Figure S3.**
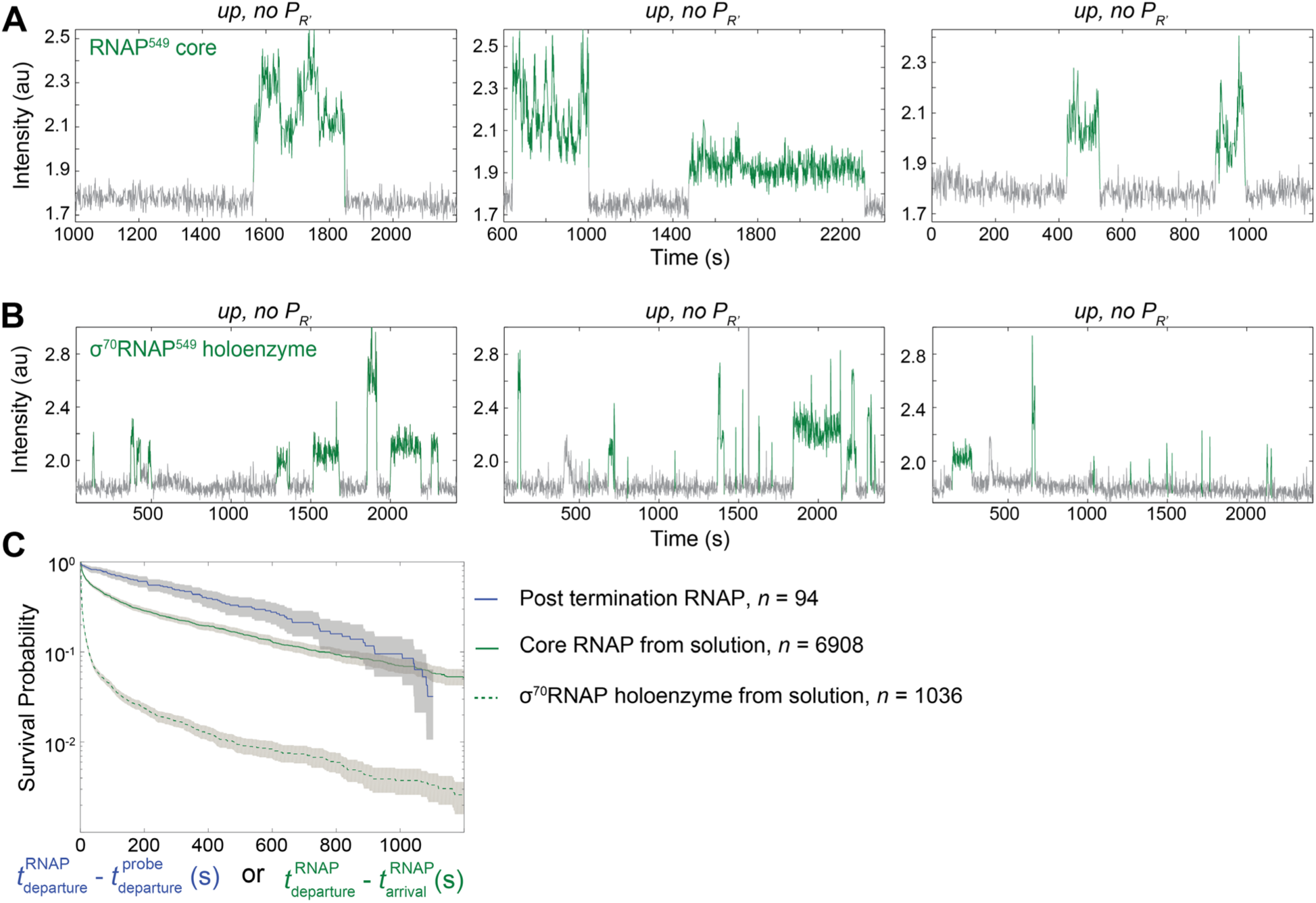
Core RNAP or σ^70^RNAP from solution binding to promoter-ablated template DNA molecules. Related to Figure 2. (**A**) Example records illustrating core RNAP binding from solution and sliding on individual template DNA molecules that lack the P_R’_ promoter sequence (*Methods*). Records are selected from an experiment in the absence of NTPs with 0.7 nM core RNAP^549^ introduced at time *t* = 0 showing RNAP^549^ emission colocalized with 3 different DNA spots. Gray color marks intervals during which no fluorescent spot was seen. (**B**) As in (A), but with 0.7 nM σ^70^RNAP^549^ holoenzyme instead of core RNAP^549^. (**C**) Survival curves of template DNA-bound RNAP^549^ (i.e., cumulative distributions of dwell times such as those shown as colored intervals in (A) and (B). Dwell times are taken from the experiments described in (A) (solid green), (B) (dashed green) and the blue subpopulation from Figure 1D, *top* (blue), which are the lifetimes of RNAP^549^ molecules retained on template DNA following termination. Shaded regions show the 90% confidence intervals of the curves determined by bootstrapping.

**Figure S4.**
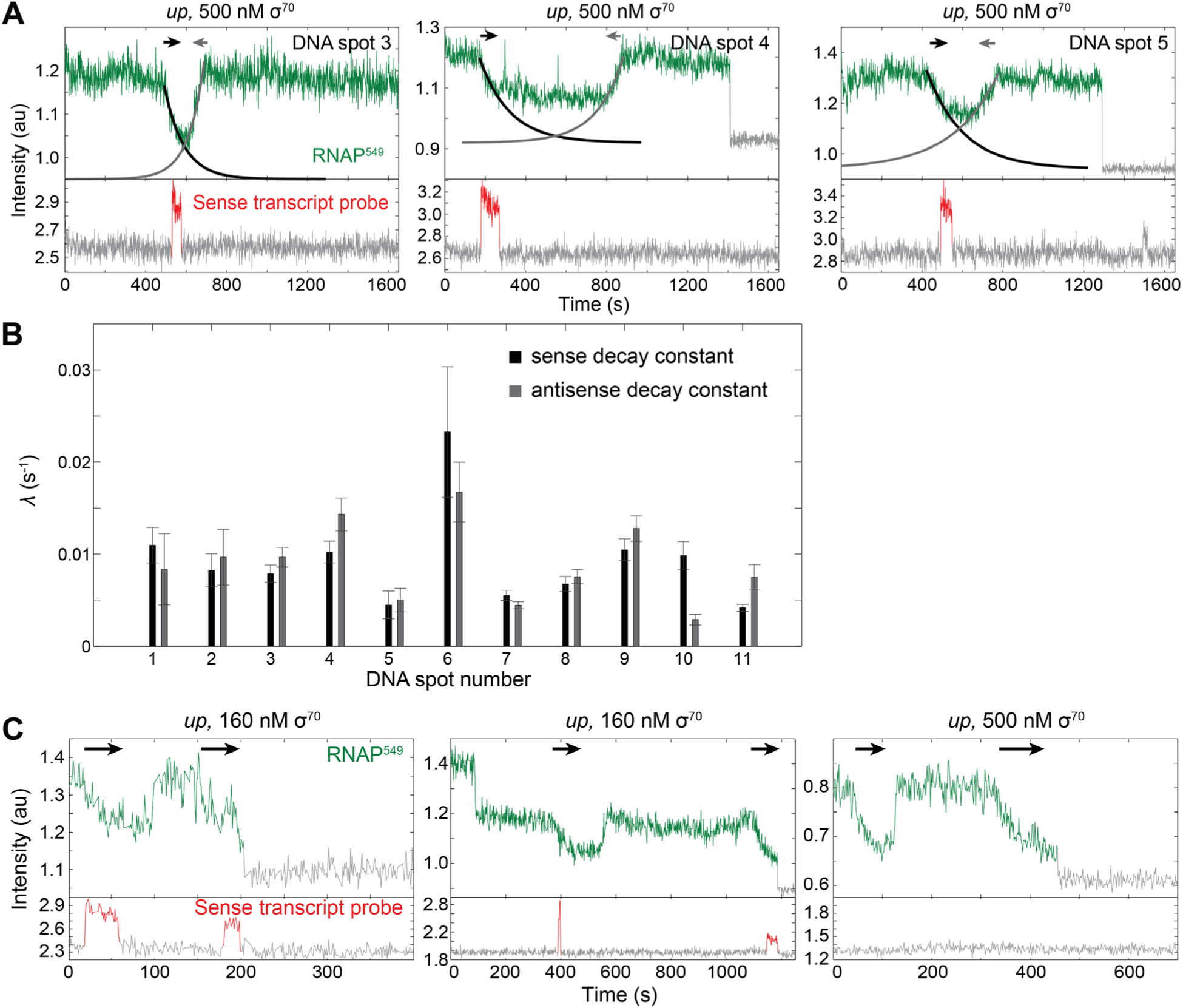
Comparing the translocation speed of single RNAP^549^ molecules during sense and antisense transcription and example single-molecule fluorescence records indicative of multiple sense transcript initiations by the same RNAP molecule. Related to Figure 3. (**A**) Single-molecule emission records, plotted as in Figure 3A, and fits to an exponential decay model (*Methods*) as in Fig S2 for the sense (black) and antisense (gray) RNAP^549^ transcription signatures. (**B**) Comparison of exponential decay constants, λ (± SE) from fit curves like those depicted in (A) drawn from 11 randomly chosen fluorescence records that exhibit both sense (black) and antisense (gray) fluorescence signatures. (**C**) Plots show records selected from two different experiments illustrating the same molecular behavior as that in Figure 3E. Each plot shows RNAP^549^ and Cy5-transcript probe emission co-localized with 3 different DNA spots. Such traces are not typical, but rather show evidence that sense transcript re-initiation following RNAP^549^ sliding may infrequently take place. Occurrences like that of the example on the right, wherein apparent steady-state elongation is observed in the RNAP^549^ TFM signal without co-localized transcript probe signal are more frequent, comprising as much as 30% of observed RNAP^549^ TFM elongation signals; we interpret these as reflecting inefficient probe hybridization due to folding of the nascent transcript. Gray color marks intervals during which no fluorescent spot was detected.

**Figure S5.**
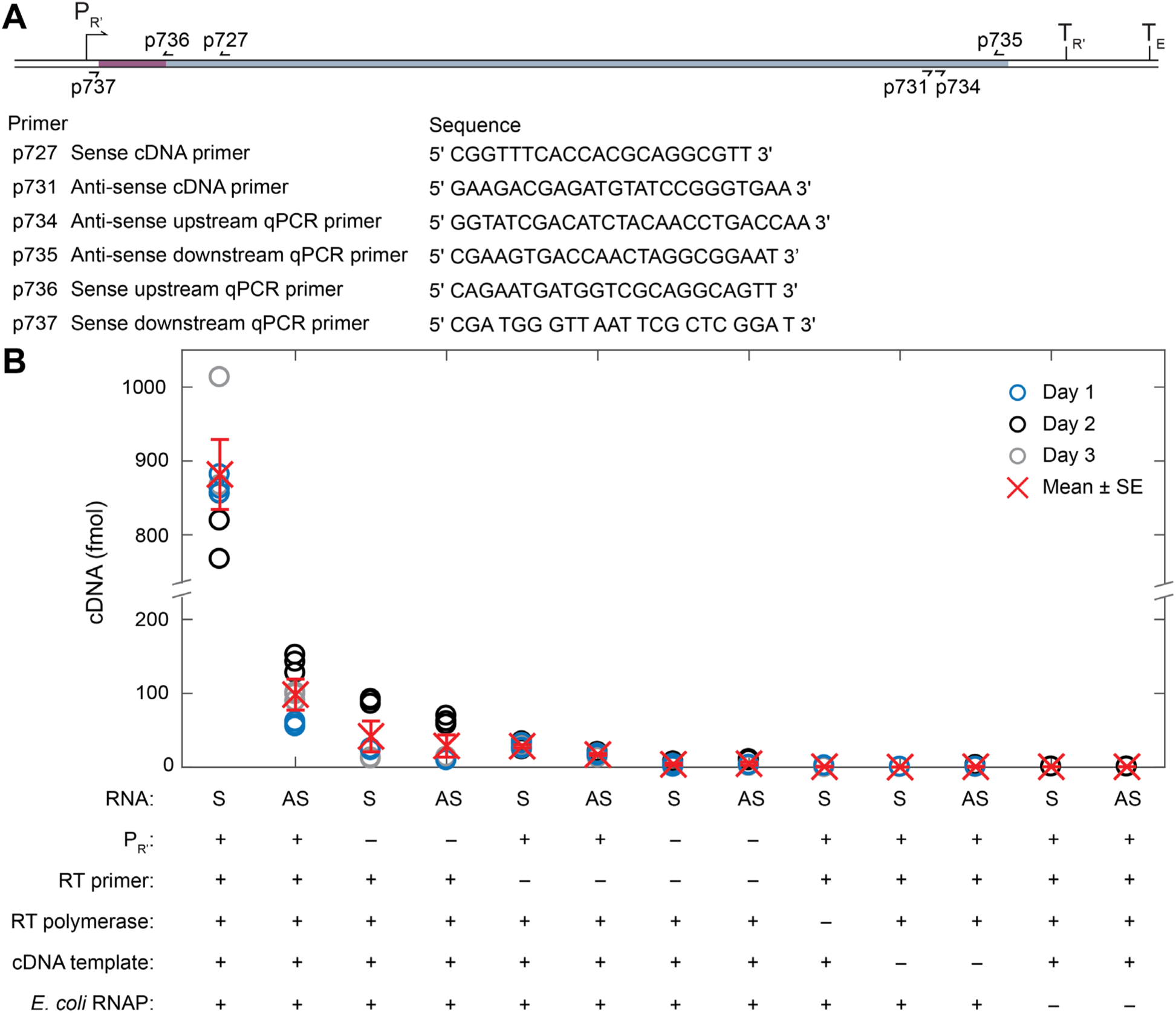
Sense and antisense transcript production *in vitro* measured by RT-qPCR. Related to Figure 4. (**A**) The transcription template and layout and sequences of the primers used for cDNA synthesis and qPCR. The template (same DNA sequence as that used in the single-molecule experiments) contains a wild type λ P_R’_ promoter region (blue, bent arrow) followed by seven tandem repeats of a 21 bp cassette (maroon), most of the *E. coli rpoB* coding region (gray) and two consecutive intrinsic terminators, λ T_R’_ and T7 T_E_, respectively. (**B**) Full results of the RT-qPCR experiments shown in Figure 4, including control experiments that withheld a selected component of the RT-qPCR assay. Colored circles indicate triplicate qPCR measurements from each of three experimental replicates conducted on different days; × indicates mean ± SE (*n* = 3). Samples designated minus P_R’_ used a version of the template in which promoter sequences were mutated (see *Methods*).

**Figure S6.**
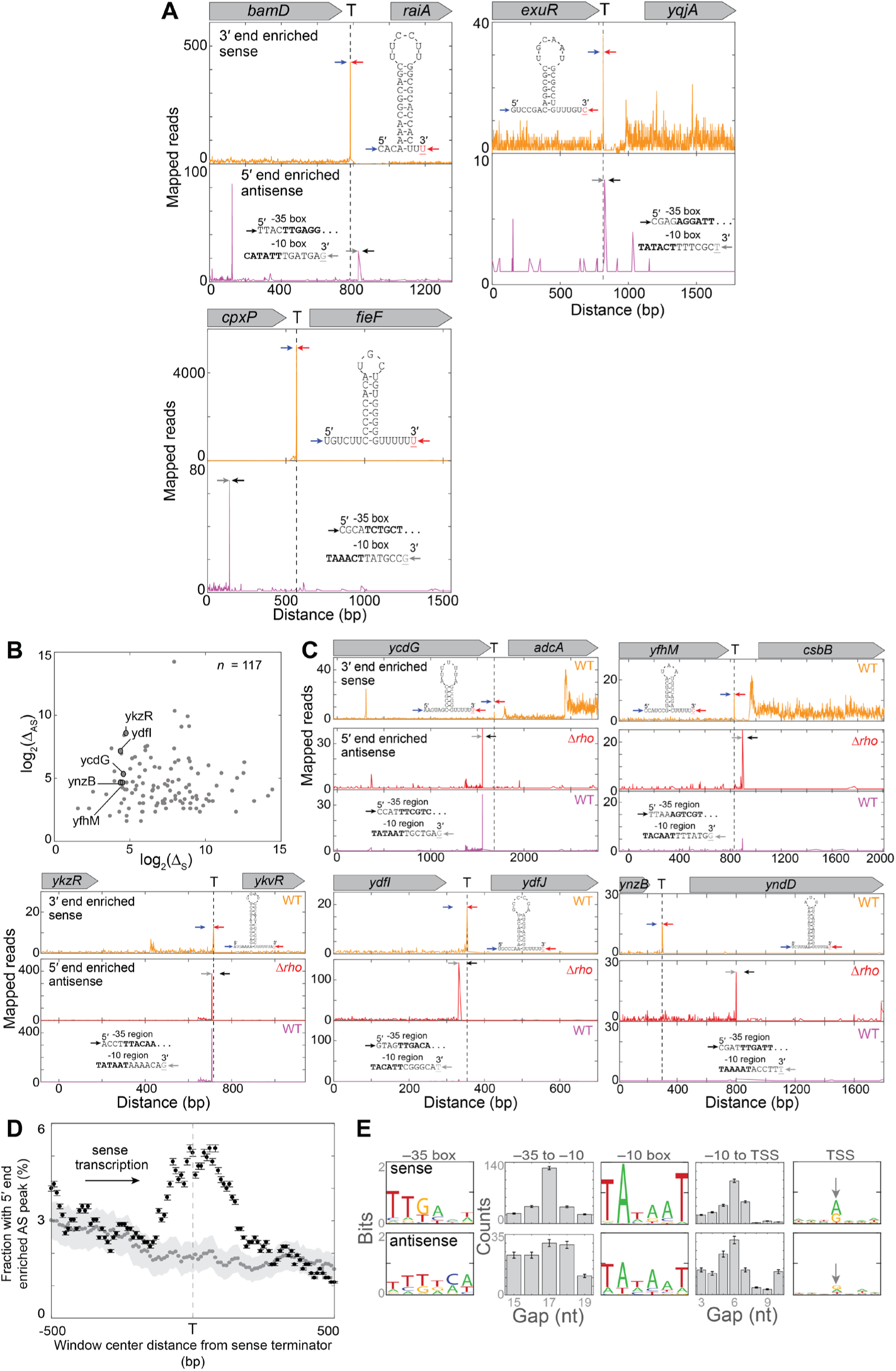
Evidence in *E. coli* and *B. subtilis* Rend-seq data for secondary initiation of antisense transcripts near positions of sense terminators. Related to Figure 5. (**A**) Data from three example *E. coli* terminators chosen from Figure 5B and plotted as in Figure 5C. (**B**) Rend-seq peak heights from a Δ*rho B. subtilis* strain, computed and plotted as in Figure 5B, for 117 of 726 terminators between genes transcribed in the same direction (see *Methods*) that show a substantial Δ_*AS*_ peak within 500 nt of the terminator Δ_*S*_ peak. (**C**) Data from five example terminators chosen from (B) and plotted as in Figure 5C. Antisense data from both wild-type (WT) and Δ*rho* strains are shown; the WT data set was normalized to have the same total reads as the Δ*rho* data set. (**D**) Antisense (AS) initiation peak frequency correlates with positions of sense terminators in the *B. subtilis* genome. Pooled data from 726 terminators between genes transcribed in the same direction (see *Methods*). Data are analyzed and plotted as in Figure 5D. Plot shows the fraction (± SE) of 200 nt-wide windows centered at the indicated distance upstream or downstream from the terminators that exhibit a peak of antisense initiation (black). Also shown is the mean ± SD of negative controls (gray) in which the same analysis was repeated 100 times using 726 randomly selected locations in the *B. subtilis* genome that lack an apparent terminator. In 100% of these 100 control replicates, the fraction at the terminator location with a 5’ end AS peak was < 2.8%, indicating that the difference between experimental data and controls was significant (*p* < 10^-2^). (**E**) Sequence consensus (illustrated as in Figure 5E) for *n* = 250 strong sense *B. subtilis* promoters (*top*) and for the *n* = 117 terminator-proximal antisense initiation sites shown in (B) (*bottom*).

